# Sex Dependent Effects of Minocycline on Contextual Fear Memory

**DOI:** 10.1101/2025.11.11.687783

**Authors:** Tzu-Ching E. Lin, Jeremy Hall, Kerrie L. Thomas

## Abstract

Recent studies have highlighted the crucial role of microglia in fear learning and memory. This supports the potential therapeutic value of the anti-inflammatory agent minocycline, which regulates microglial activity, in psychiatric disorders involving dysregulated emotional memory. To assess the impact of minocycline during a tractable intervention window, we examined its acute systemic effects on the reconsolidation, extinction, and forgetting of contextual fear memory (CFM) in rats. Our findings show that the effects of minocycline on fear memory and anxiety-like behaviours depend on sex, memory activation (i.e., whether it was retrieved), and memory age. In males with recently acquired CFM, minocycline given before retrieval enhanced extinction and reduced fear expression. When administered without retrieval, it disrupted passive forgetting, increasing evoked fear behaviour. This extinction-enhancing effect did not occur with older (28-day) memories rather it strengthened CFM through enhancing reconsolidation. In contrast, in females, minocycline increased the expression of recent fear memories regardless of retrieval, without affecting extinction or forgetting. This elevated fear expression appeared linked to heightened anxiety-like behaviour. These findings suggest that minocycline, or other microglia-targeting agents, would need to be used in a highly selective and targeted manner to therapeutically modulate pathological fear memories. Minocycline may be beneficial as an adjunct to exposure therapy in men explicitly retrieving recent fear or trauma-related memories, but outside these conditions, it may worsen fear expression. These results help explain the mixed findings from therapeutic trials using minocycline in psychiatry and support the application of a more precision approach in any future applications.

## Introduction

Dysregulated fear memory formation and processing is a key feature of many psychiatric disorders [1, 2], including post-traumatic stress disorder (PTSD)[3] , anxiety disorders [4, 5], depression[6, 7], and schizophrenia [8–10]. The modification of fear memory formation and processing thus represents an important target for the development of novel therapies for psychiatric conditions.

Rodent studies suggest microglia contribute to hippocampal-dependent fear learning and memory. Microglia play key roles in physiological processes such as synaptic maturation and plasticity, which support learning and memory, in both the developing and adult brain [11, 12]. Indeed, mice lacking the CX3CR1 receptor for the chemokine CX3CL1 (fractalkine) specific to microglia show impairments in hippocampal plasticity, learning, and neurogenesis [13, 14]. A role for microglial-derived BDNF in hippocampal plasticity and learning has also been suggested [15, 16]. Additionally, there is increasing evidence that microglia play a role in the pathogenesis of psychiatric disorders [17–20]. An important study showed that adolescents treated with minocycline, a second-generation tetracycline derivative with anti-inflammatory and immune-modulatory effects [21–24] had lower subsequent risk of psychosis [25] indicating that inhibiting microglial activation can alter risk for psychotic conditions. Gerst et al. have also reported that minocycline can reduce symptoms of PTSD [26], and trials are ongoing in individuals with depression and raised inflammatory markers. Drugs that target microglia activation, such as minocycline, have thus attracted considerable interest as potential therapeutic agents in psychiatry, although negative results from trials in psychosis and depression have also been reported [27–30].

There is increasing evidence that microglia may play a particular role in post-learning modifications of fear memories. Depleting microglia or inhibiting their activity with minocycline has been shown to prevent the forgetting, or passive loss, of fear memory[31]. It has also been reported that minocycline can facilitate the extinction of fear memories [32]— considered a new learning process initiated by the retrieval of a non-reinforced memory where stimuli gain additional meaning resulting in the loss of memory associated behaviours [33–35]. These data indicate that there may be contrasting roles for microglia in specific fear memory processes, facilitating passive forgetting in the absence of memory retrieval but inhibiting extinction when memory is actively retrieved. Less is currently known about any potential role of microglia in reconsolidation, the process through which stable consolidated memories can enter a transient labile state allowing the memory to be updated or strengthened[36]. The involvement of microglia in post-learning modification of fear memories may thus represent an opportunity for therapeutic intervention.

Recent studies show that acute minocycline administration can attenuate fear memory consolidation during learning in humans[37]. However, the effects of minocycline on post-consolidation memory processes remains to be full characterised. The potential effects of minocycline on post-consolidation fear memory are particularly important to assess, given that it is this stage of memory processing which is most likely to be accessible clinically in conditions such as PTSD. Critically however any therapeutic use of compounds like minocycline is likely to depend critically on the conditions under which memory is retrieved, and whether these trigger reconsolidation (updating and strengthening memory) or extinction (weaking memory expression) [38]. Here, we therefore examine systemic minocycline’s effects on the reconsolidation, and extinction of contextual fear memory (CFM), as well as on forgetting (passive memory loss) [39]. Given the known sex differences in the presentation of a range of fear-related psychiatric conditions, and potential sex-specific effects of minocycline [40], we also explicitly examined the effects of minocycline in male and female rats.

## Materials and Methods

### Animals

A total of 152 male and 88 female Lister Hooded rats (Envigo, UK) were used in this study. The rats were held in light/dark reversal holding room maintained at 21°C, lights out from 10 a.m. to 8 p.m. The rats were approximately 3 months old at the start of the experiment and were housed in same-sex pairs in large rat home cages in the holding room. Behavioural training began at approximately 10.30 a.m. on each day. Animal had free access to food and water throughout the whole experiment. All procedures were conducted in accordance with local Cardiff University Ethical Committee approval and the United Kingdom 1986 Animals (Scientific Procedures) Act (Project license P0EA855DA).

### Behavioural paradigms

A paradigm using a single footshock exposure in a novel context was used. During 3-min contextual fear conditioning (CFC) training trial, each animal received a single scrambled footshock (unconditioned stimulus (US), either 0.35mA, 0.5mA or 0.7mA for 2s) after being placed in one of two randomly assigned conditioning contexts (context conditioned stimulus, CS) for 2 min and left in the conditioning chamber (Med Associates Inc., Vermont, USA) for a further 1 min. Animals were returned to home cage after conditioning. For each trial, pairs of animals were transferred between the home cages and behavioural testing room in the same transport box. Contextual fear memory was tested 48 hours later by measuring the animal’s conditioned freezing response (immobility except for respiration) upon return to the conditioned chamber for 2 min (long term memory test, LTM). Additional 2 min LTM tests were given at the times indicated in the figures. Cohorts of animals underwent extinction training (Extinction) 48 hrs after CFC by the unreinforced (no US) re-exposure to conditioned chamber for 10 min. To determine the presence of extinction, a reminder CS was presented at the end of a 2 min context re-exposure (Reminder) and extinction identified by an increase in freezing response at a subsequent LTM [41]. During the Reminder session, animals were returned to the conditioning chamber for 2 min and a single 2 s 0.25mA scrambled foosthock was delivered at the end of the session. This level of footshock is not strong enough to establish CFM but is sufficient to reveal freezing behaviour previously extinguished CFM during a later LTM [42]. Freezing behaviour was used as measure of fear. One unit of freezing response was defined as the continuous absence of movement other than respiratory motion in 1 s sampled every 10s. Freezing responses were electronically recorded using IR cameras (JSP Electronics Ltd, China) during each trial and quantified ofline as the percentage of time spent freezing per 2 min epoch (except the post US epoch during CFC, 1 min) by an observer blind to the experimental group.

The present study used different intensities of footshock in males and females to assess the impact of minocycline on modulating the strength of CFM. We initially used a 0.5mA intensity of footshock to establish CFM in the current study. However, our observations showed female rats tended to display lower levels of freezing behaviour than the males when they were subjected to a similar US during conditioning which is concordant with previous observations [43, 44, Supplementary Fig. 1] and when using a 0.5mA US the behavioural expression of CFM was short-lived in females (Figure 3C,D). Therefore, 0.7mA footshock was used to establish a reliable and enduring CFM in the female rats.

### Minocycline injection

Minocycline hydrochloride (Merck, UK, Cat#: M9511) was freshly dissolved in sterile 0.01M PBS and 1M NaOH prior use. The pH of the final working solution was at approximately 5.5. A single intra-peritoneal injection of minocycline (40mg/kg) or vehicle (vol) was given 30 min prior to the designated training or LTM session, or for No Recall groups at the same equivalent times post CFC. The dose and injection route was chosen based on literature reviews [32, 45].

### Statistical Analysis

Group sizes were defined by prior power analysis based on estimates based on our previously published data using similar behavioural paradigms. Results were analyzed with two-way ANOVA with repeated measures (factors included: minocycline treatment, Test (2min recall and LTM tests, extinction and reminder sessions) or US intensity, and independent samples t-test. Because experiments on males and females were not run concurrently, sex was not included as a factor in formal analysis. Significance was accepted when *p* < 0.05

## Results

### Minocycline affects the expression of contextual fear memory in a retrieval and sex-dependent manner

We investigated the effect of minocycline on forgetting and the reconsolidation and extinction of conditioned fear memory (CFM) in male and female rats. Briefly, two days after contextual fear conditioning (CFC), cohorts of rats received a single systemic injection of either minocycline or phosphate buffered saline (PBS) 30 mins prior to exposure to the conditioned context for either 2 min (2 min Recall group) or 10 min (10 min Extinction group) (Fig. 1A). We have previously shown that 10 min non-reinforced exposure to the training context engages the extinction of conditioned freezing after CFC [46], whereas a 2 min exposure does not, preferentially initiating reconsolidation[35, but see 42]. Animals in a third No Recall group only received minocycline or PBS injections and were returned to their home cages thereafter without re-exposure to the conditioned context. This group served to measure the effect of minocycline on the forgetting of established CFM when the fear memory was not reactivated. Subsequently all animals were returned to the conditioned chamber 5 days (LTM1) and 26 days (LTM2) later to assess the effect of minocycline on CFM and extinction. Because females had weaker CFM using a 0.5mA US during CFC (see Materials and Methods, Behavioural Paradigms), we used 0.7mA to match male freezing behaviour and test minocycline’s effects on similar levels of CFM.

**Fig. 1.**
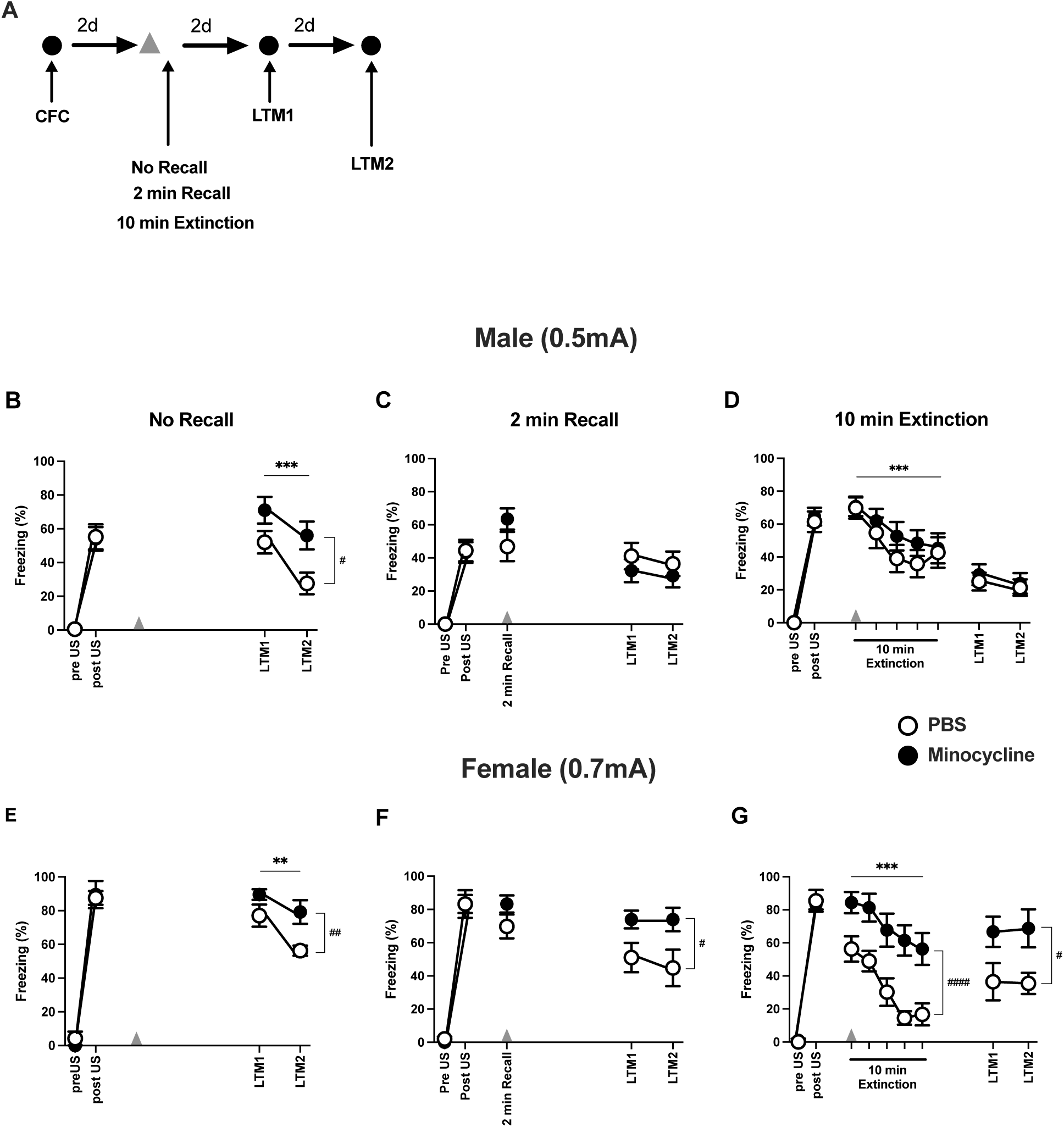
Minocycline affects the expression of contextual fear memory in a sex dependent manner. **A**: Schematic of the CFC (contextual fear conditioning) and contextual fear memory (CFM) behavioural protocol. CFC was established using a single 2s scrambled footshock US (males 0.5 mV, females 0.7 mV) given 2min into a 3 min exposure to a novel context. Arrowhead shows the administration of minocycline (40mg/kg, i.p.) or PBS 30 min prior to either a 2min CFM recall trial (2 min Recall), a 10 min Extinction trial (10 min Extinction, 5 x 2min epochs, 1-5), or at an equivalent time without recall (No Recall). Rats were exposed to the conditioning context for 2 min 2d (long-term memory test 1, LTM1) and 23d (LTM2) after either a minocycline or control injection. **B-D The effect of minocycline in male rats**. **(B)** No Recall: There was a decline in conditioned freezing between the two LTM tests with minocycline administered males showing greater freezing behaviour than those injected with PBS (Test: F_(1, 30)_ = 18.713, p < 0.001; Mino: F_(1, 30)_ = 6.518, p = 0.016; Test x Mino: F_(1, 30)_ = 1.050, p = 0.314). **(C)** 2 min Recall. There was no difference between minocycline and PBS-injected rats at CFM Recall (t (30) = 1.527, p = 0.137) or during LTM tests (Test: F_(1, 30)_ = 1.136, p = 0.295; Mino: F_(1, 30)_ = 0.689, p = 0.413; Test x Mino: F_(1, 30)_ = 0.045, p = 0.833). **(D)** 10 min Extinction: During the 10 min extinction session, both minocycline and PBS-injected rats showed reduced freezing behaviour across the 10 min extinction training (Extinction: F_(2.871, 86.123)_ =9.785, p < 0.001, Mino: F_(1, 30)_ = 0.640, p = 0.430, Extinction x Mino: F_(2.871, 86.123)_ =0.542, p = 0.647). Both treatment groups also showed equivalent low levels of freezing at LTM 1 and LTM 2 after extinction training (Test: F_(1, 30)_ =1.676, p = 0.205: Mino: F_(1, 30)_ = 0.206, p = 0.653; Test x Mino: F_(1, 30)_ = 0.052, p = 0.821). **E-G The effects of minocycline in female rats**. **(E)** No Recall: female rats showed decreased freezing behaviour over the two LTM tests, however minocycline-treated rats showed greater freezing behaviour than the PBS-injected animals (Test: F_(1, 14)_ = 10.714, p =0.006, Mino: F_(1, 14)_ = 9.409, p = 0.008, Test x Mino: F_(1, 14)_ =1.190, p = 0.294). **(F)** 2 min Recall: There was no effect of minocycline on freezing behaviour at CFM recall 30min after the injection (t_(14)_ =1.522, p = 0.15). However, minocycline-injected rats showed greater conditioned freezing responses compared to PBS controls at LTM1 and LTM2 tests (Tests: F_(1, 14)_ = 0.273, p = 0.610: Mino: F_(1, 14)_ = 6.559, p = 0 .023; Test x Mino : F_(1, 14)_ = 0.273, p = 0.610). **(G)** 10 min Extinction: During the 10 min extinction training session, all rats showed decreases of freezing response within the 10 min extinction session with minocycline-injected females displaying greater freezing responses than PBS-injected rats (Extinction: F_(4, 56)_ =16.835, p < 0.001; Mino: F_(1, 14)_ =17.362, p < 0.001; Extinction x Mino: F_(4, 56)_ =0.907, p = 0.466). During the LTM tests, minocycline-injected animals also showed greater fear responses compared to vehicle controls (Test: F_(1, 14)_ = 0.008, p = 0.930; Mino: F_(1, 14)_ = 6.367, p = 0.024, Test x Mino: F_(1, 14)_ = 0.071, p = 0.793). Results are the Mean ± SEM. Per group, n = 16 for males (B - D) and n = 8 for females (E - G). * p < 0.05, ** p < 0.01, *** p < 0.001 compared between LTM tests; # < 0.05, ## p < 0.01 compared between groups.

The results showed that a single acute administration of minocycline modulated the expression of CFM, but its effect differed by sex. In male rats, minocycline treatment given in the absence of fear memory retrieval (No Recall group) increased freezing behaviour measured at subsequent memory tests, LTM1 and LTM2 (Fig. 1B). However, there was no increase at LTM1 and LTM2 when minocycline was given either prior to a retrieval session that might promote the reconsolidation (2 min Recall, Fig. 1C), or extinction of CFM (10min Extinction, Fig. 1D). In addition, there was no acute effect of minocycline on the freezing behaviour during the 2 min CFM retrieval session (Fig, 1C), or on the diminution of freezing behaviour during a 10 min retrieval session (within-session extinction, Fig. 1D). Thus in males, minocycline exerted a differential effect on long-term fear memory expression depending upon whether memory was retrieved or not.

In females, minocycline increased freezing behaviour at LTM1 and LTM2 when administered in the absence of retrieval (No Recall, Fig. 1E) as well as before of a brief (Fig. 1F) or prolonged re exposure the training context which induced extinction (Fig. 1G). Minocycline did not affect the ability of females to undergo extinction with 10 min context re-exposure as indexed by a reduction in conditioned freezing responses despite higher responses initially. Therefore, in females minocycline resulted in a general and persistent increase in freezing behaviour independent of whether memory was not retrieved, retrieved or retrieved and extinguished after administration.

The enhancement of conditioned freezing by minocycline at recent (2 or 4 days after CFC, LTM1) or more remote CFM tests (26 days after CFC, LTM2) cannot be explained by a systematic difference in CFM acquisition during conditioning because prior to administration, PBS and minocycline groups showed equivalent levels of responding to the US during the CFC training (Supplementary Table 1). These results suggest that minocycline modulates the expression of an established fear memory in a retrieval and sex-dependent manner: 1)

Without CFM retrieval, acute minocycline enhances the expression of fear memory in both females and males; 2) when given prior to CFM retrieval, minocycline enhances the expression of CFM in female rats only.

### Minocycline enhances extinction in males

We observed that the effect of minocycline on fear memory in males depended on whether the memory was retrieved or not. Our results are consistent with minocycline triggering extinction in the 2 min Recall group in males, a process that usually requires a longer unreinforced context exposure as observed in the 10 min Extinction group. To test the hypothesis that minocycline might augment extinction, we trained males as well as females to show a robust CFM using the higher intensity 0.7 mA footshock which produces a stronger contextual fear memory (CFM) [44, 47, 48] (Supplementary Fig. 1) known to be more resistant to extinction [49]. Subsequently, both males and females received two 10-minute extinction sessions (Extinction 1 and Extinction 2), spaced at least 21 days apart, followed by a low-intensity 0.25 mA reminder footshock. Spontaneous recovery of freezing over time, or reinstatement of fear after the reminder, would evidence extinction rather than a loss of CFM [50].

Male rats given minocycline or PBS showed similar reductions in freezing behaviour during the first extinction session, and also at LTM1 and LTM2 with no evidence of spontaneous recovery of the freezing response at the second LTM2 test 21 days later (Fig. 2A). In marked contrast, conditioned behavioural responses were significantly lower during Extinction 2 and after the second extinction session (LTM3 and LTM4) in minocycline injected males. The reminder US increased conditioned freezing at LTM5 in both groups commensurate with minocycline modulating extinction rather than the original CFM.

**Fig. 2.**
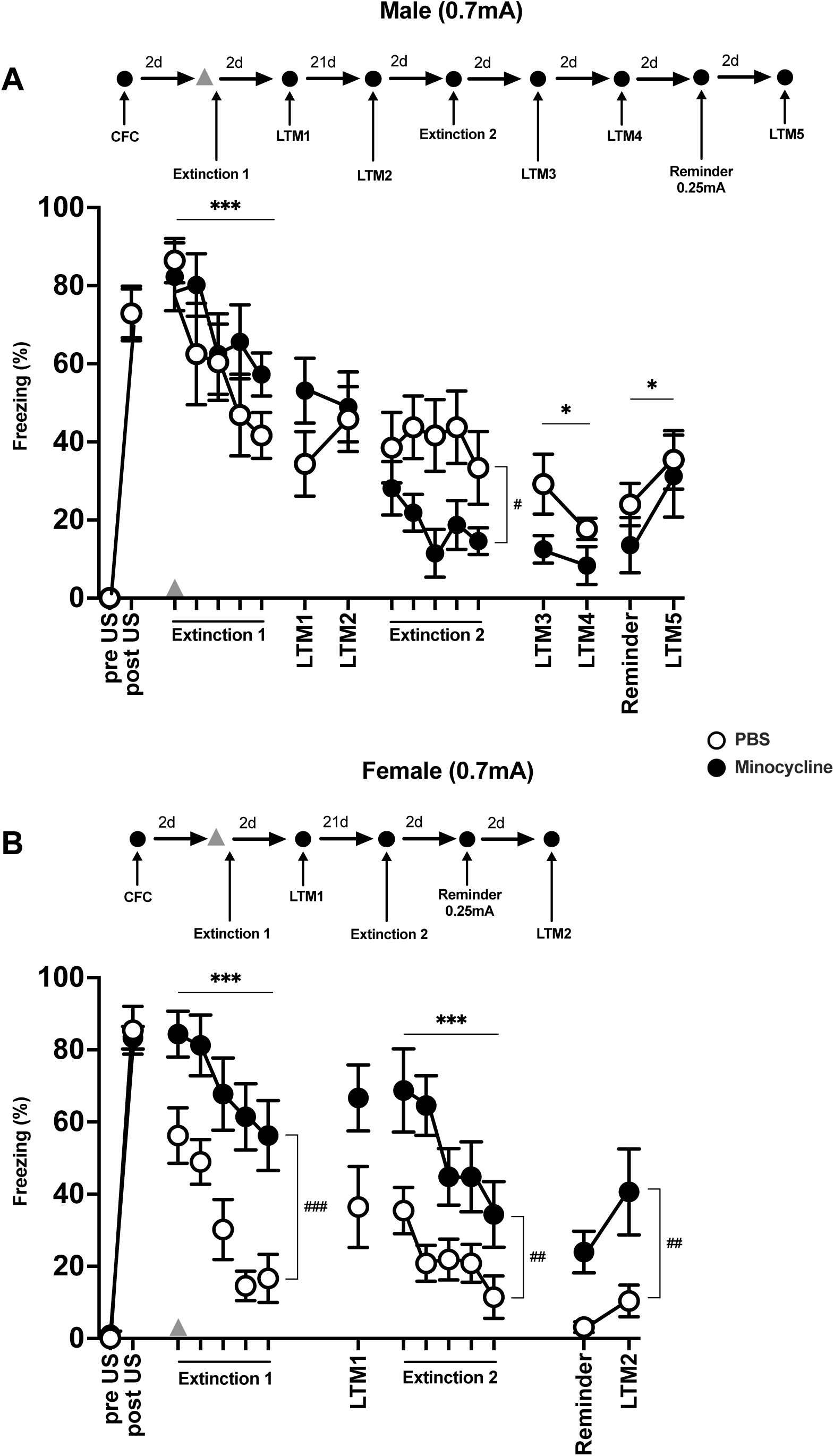
Minocycline enhances extinction in males only. **(A) The effect of minocycline on extinction of robust CFM with two extended trials in male rats**. Top: Schematic of the CFC and CFM behavioural protocol. Males were conditioned using a 2s, 0.7mV US. Minocycline (40mg/kg, i.p.) or PBS (arrowhead) was given 30 min prior the first of two 10 min extinction trials (Extinction 1 and 2). LTM denotes 2 min CFM tests. Reminder: a 2 s 0.25mA US was given at the end a 2 min LTM test to evoke recovery of CFM after extinction. **(A) The effect of minocycline on extinction with two extended trials in male rats**: Both minocycline and PBS-treated males showed a reduction in conditioned freezing behaviour during the first extinction training session (Extinction 1: F_(4, 56)_ = 9.021, p < 0.001, Mino: F_(1, 14)_ = 1.046, p = 0.324, Extinction 1 x Mino: F_(4, 56)_ = 1.305, p = 0.279) and showed no further reduction of freezing behaviour between subsequent LTM1 and LTM2 tests (Test: F_(1, 14)_ = 0.565, p = 0.465, Mino: F_(1, 14)_ = 0.996, p = 0.335, Test x Mino: F_(1, 14)_ = 2.595, p = 0.130). However, minocycline administered males showed lower freezing responses compared to PBS males during Extinction 2 (Extinction 2: F_(4, 56)_ = 0.862, p = 0.492, Mino: F_(1, 14)_ = 9.244, p = 0.009, Extinction 2 x Mino: F_(4, 56)_ = 0.692, p = 0.601). Fear memory responses at subsequent tests (LTM3, LTM4) further reduced in both groups (Test: F_(1, 14)_ = 4.937, p = 0.043, Mino: F_(1, 14)_ = 4.345, p = 0.056, Test x Mino: F_(1, 14)_ = 1.075, p = 0.317). Nevertheless, in both groups the reminder US increased freezing behaviour at LTM5 compared to that measured during the Reminder session before US delivery (Reminder: F_(1, 14)_ = 7.240, p = 0.018, Mino: F_(1, 14)_ = 0.563, p = 0.466, Reminder x Mino: F_(1, 14)_ = 0.332, p = 0.573). **(B) The effect of minocycline on extinction of robust CFM with two extended trials in female rats**: Top: schematic of behavioural protocol. Female rats were CFC using a 2s, 0.7mV US. Minocycline-treated females showed greater levels of freezing behaviour compared to the PBS females despite both groups showing a decline in freezing behaviour across the first 10 min extinction trial (Extinction 1: F_(4, 56)_ =16.835, p < 0.001, Mino: F_(1, 14)_ =17.362, p < 0.001, Extinction x Mino F_(4, 56)_ =0.907, p = 0.466), and marginally higher freezing behaviours during LTM1 (t(14) = 2.081, p = 0.056). Both groups showed further extinction of freezing behaviour during a second extinction session, again with the minocycline-treated group showing higher fear responses than the PBS group (Extinction 2: F_(4, 56)_ =7.387, p < 0.001, Mino: F_(1, 14)_ =12.825, p = 0.003, Extinction 2 x Mino: F_(4, 56)_ = 1.265, p = 0.295). The minocycline group showed higher fear responses at subsequent reminder and test sessions, with the reminder evoking a trend towards an increase in fear responses at LTM2 (Reminder: F_(1, 14)_ = 4.101, p = 0.062, Mino: F_(1, 14)_ = 10.229, p = 0.006, Reminder x Mino: F_(1, 14)_ = 0.628, p = 0.441). Results are the mean ± SEM. n = 8 per group. * p < 0.05, ** p < 0.01, *** p < 0.001 compared between LTM tests; # < 0.05, ## p < 0.01 compared between groups.

**Fig. 3.**
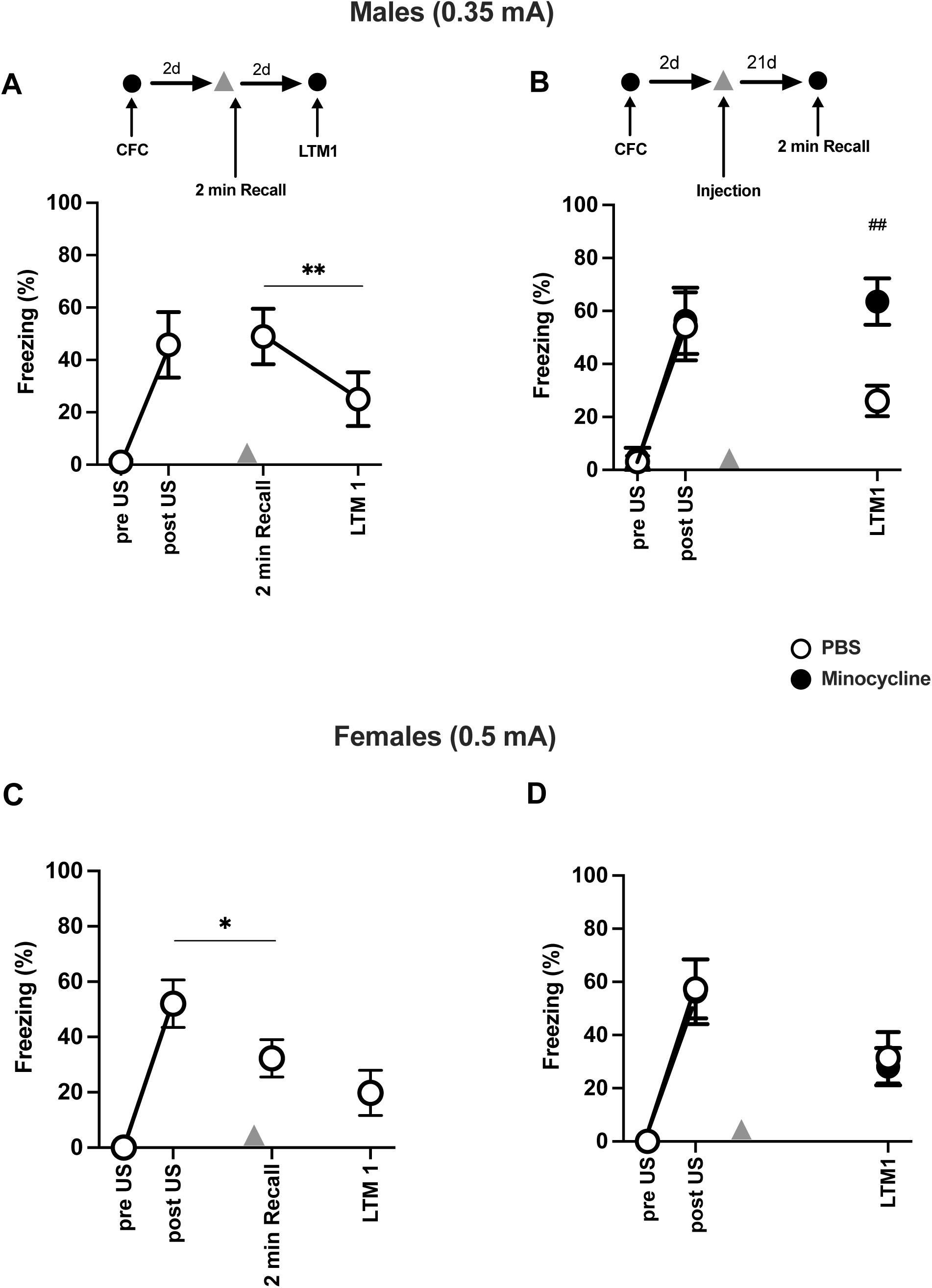
Minocycline without memory retrieval strengthens weak fear memory in males only. Tops: Schematics of CFC and CFM (a 2 min test, LTM) behavioural protocols. CFC was established with a 2s 0.35mA or 2s, 0.5mA US in males and females, respectively. Arrowhead: Intraperitoneal injection of minocycline (40mg/kg) or phosphate buffer saline (PBS) 30 min either before a 2 min recall session (2 min Recall) or at an equivalent time after CFC with no recall. **(A) Males:** CFM established with a 0.35mV US was weak and declined over 4 days (*t(*7) = 3.643, p = 0.008). **(B)** Male rats that were administered minocycline without recall showed greater levels of conditioned freezing behaviour than those treated with PBS at the LTM1 test 23d later (*t*(14) = – 3.575, p = 0.003). **(C) Females**: CFM established with 0.5mA US declined quickly between Post US and 2 min recall test (*t*(7) = 2.569, *p* = 0.037) and low conditioned fear was maintained at LTM1 (*t*(7) = 1.122, *p* = 0.299). **(D)** Female rats administered minocycline after CFC in the absence of CFM recall showed equivalent low levels of conditioned fear behaviour to PBS administered females 23 days after CFC (*t*(14)= 0.610, p = 0.552). Results are the mean ± SEM. n = 8 per group. * p < 0.05, ** p < 0.01 compared between LTM tests; # < 0.05, ## p < 0.01 compared between groups.

As similarly seen previously, female rats administered minocycline prior to extinction training, showed both within-session reductions in freezing behaviour during extinction (Extinction1), which were maintained at LTM1 as evidence of extinction despite showing higher levels of conditioned freezing behaviour than PBS-treated rats (Fig. 2B). There was no evidence of recovery of the CFM in the first 2 min of the second extinction session 21 days later compared to LTM1 in both groups. Both groups also showed further extinction of conditioned fear responses with a second 10 min extinction training session 21 days later, again with minocycline-treated females showing higher responses than their PBS counterparts. The difference in freezing behaviour between the two groups was maintained during the reminder and subsequent test session (LTM2) with a trend for the US reinstatement of freezing behaviour at LTM2. This effect was most marked in the minocycline group as evidence of intact extinction.

These results show that minocycline given prior to extinction training selectively augments the extinction of CFM in males but not in females.

### Minocycline in the absence of retrieval strengthens fear memory in males

Given that both male and female rats showed increased conditioned fear responses in the No Recall group, we hypothesized that acute systemic minocycline administration without the retrieval of CFM may strengthen an established fear memory. To assess this possibility, rats were conditioned using a lower intensity footshock prior to receiving minocycline or PBS injections 2 days later in the absence of CFM retrieval (Fig. 3). We used a 0.35mA footshock to establish CFC in the male rats. CFM established with this level of footshock was weak such that conditioned freezing 2 days post conditioning had diminished by 4 days (Fig. 3A). In a separate cohort, when minocycline was given 2 days after CFC training in the absence of CFM retrieval, minocycline-treated males showed a significantly greater level of conditioned fear than PBS-treated animals when CFM was probed remotely 21 days later (Fig. 3B).

In female rats, using a 0.5mA intensity of footshock during CFC resulted in a weak and short-lasting CFM as shown by diminished conditioned freezing responses during testing by 4 days later (Fig. 3C). Minocycline administered 2 days after CFC with a 0.5mA US without CFM retrieval had no effect on the levels of conditioned freezing behaviour measured 21 days later (Fig. 3D).

Thus, while CFC using a 0.35mA or 0.5mA US in males and females respectively produced a similarly weak and short-lived CFM as measured 4-23 days later, administering minocycline without CFM retrieval strengthened CFM in males but not in females.

### Minocycline enhances the reconsolidation of remotely established memories in males

To investigate the effects of minocycline on the reconsolidation and extinction of a long-established (remote) fear memory, rats were given minocycline or PBS injections 30 min prior to retrieval 24 days after CFC training.

In males, the standard 0.5mA US produced equivalent levels of freezing behaviour measured at LTM1 and 21d after CFC in the minocycline- and PBS-injected groups establishing that both groups showed similar level of remote CFM prior to minocycline administration before a second test 2 days later (2 min Recall, Fig. 4B). Minocycline injection did not affect the immediate expression of CFM at 2 min Recall. However, the minocycline treated group showed significantly *greater* levels of freezing responses at a retrieval test 2 days later (LTM2), during extinction training, despite showing within session decreases in freezing behaviour, and during a subsequent recall test 2 days later (LTM3). Since the levels of conditioned freezing were already very low for the PBS treated animals at LTM3, precluding the value of a further extinction training session, we gave a low 0.25mA US reminder to probe CFM and extinction. The reminder was effective in evoking an increase of conditioned fear behaviour measured at LTM4 in the PBS group but not the minocycline group. The failure to reinstate freezing behaviour by the reminder indicates that minocycline targeted the long-established CFM rather than extinction processes that might be engaged following retrieval at 2 min Recall. In males the higher freezing responses before, during and after extinction training are consistent with minocycline enhancing the reconsolidation of remote CFM.

**Fig. 4.**
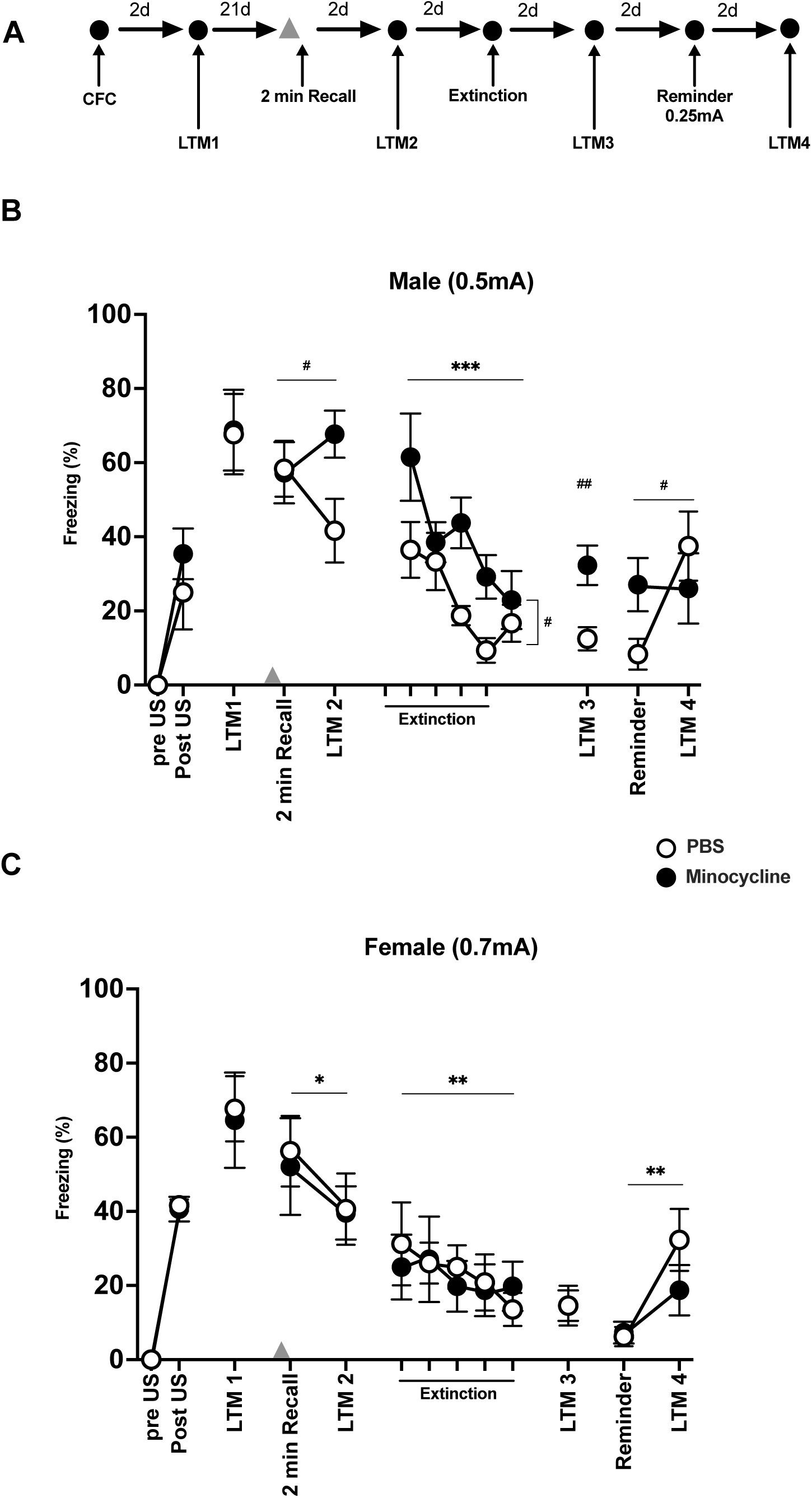
Minocycline treatment strengthens remote fear memory in males only. **(A):** Schematics of the CFC and CFM (LTM tests) behavioural protocols. Males and females were conditioned using a 2s, 0.5mV or 2, 0.7mV US respectively. Minocycline (40mg/kg, i.p.) or PBS (arrowhead) was given 30 min prior a 2min recall session 21 days after the LTM1 test. Extinction: a 10 min non-reinforced re-exposure to the CFC context. Reminder: 2 s 0.25mA US given at the end a 2 min LTM test to evoke recovery of CFM after extinction. **(B) Males**: Rats in both minocycline and PBS groups showed equivalent levels of conditioned freezing prior at LTM1 to minocycline injection (*t*(14) = 0.068, p = 0.947). There was an interaction between test (2 min recall and LTM 2) and treatment (Test: F _(1,14)_ = 0.453, p = 0.512, Mino: F_(1, 14)_ = 1.585, p = 0.229, Test x Mino: F_(1, 14)_ = 8.511, p = 0.011) such that lower freezing behaviour was observed in PBS compared to minocycline treated rats at LTM2. During extinction, there was a decline of freezing behaviours in both groups, however, minocycline injected rats showed higher levels of freezing compared to PBS rats (Extinction: F _(4,56)_ = 8.774, p < 0.001, Mino: F_(1, 14)_ = 6.882, p = 0.020, Extinction x Mino: F_(4, 56)_ = 1.393, p = 0.248). The minocycline group showed greater levels of freezing behaviour at LTM 3, (t( 14) = 3.199, p = 0.006).The reminder stimulus caused some recovery of freezing behaviour in the PBS group but not the minocycline group at LTM4 (Reminder: F_(1, 14)_ = 5.458, p = 0.035, Mino: F_(1, 14)_ = 0.153, p = 0.701, Reminder x Mino: F_(1, 14)_ = 6.296, p = 0.025). **(C) Females**: Comparable levels of conditioned fear were seen at LTM1 in both groups before treatment (*t* _(14)_ = 0.200, p = 0.844). After 21 days, both minocycline and PBS-treated female rats showed similar levels of conditioned freezing behaviour despite a decline in responses between the 2 min recall and LTM2 test (Test: F _(1,14)_ = 5.191, p = 0.039, Mino: F_(1, 14)_ = 0.041, p = 0.842, Test x Mino: F_(1, 14)_ = 0.064, p = 0.804). Freezing behaviour during extinction was markedly low with no decline during the session and no group differences (Extinction: F_(4,56)_ = 1.298, p = 0.282, Mino: F_(1, 14)_ = 0.023, p = 0.882, Extinction x Mino: F_(1, 14)_ = 0.374, p = 0.826), and no group differences were seen at LTM3 (t (14) = 0.000, p = 1.000. Comparison of freezing at LTM2 and LTM 3 indicated a reduction of freezing behaviour by LTM3 in both groups as evidence of extinction by extended context exposure (Test: F_(1, 14)_ = 21.248, p < 0.001, Mino: F_(1, 14)_ = 0.004, p = 0.949, Test x Mino: F_(1, 14)_ = 0.009, p = 0.926). The reminder US evoked an increase in freezing behaviour in both the PBS and minocycline-treated rats at LTM 5 compared to responses in the Reminder session (Reminder: F_(1, 14)_ = 11.484, p = 0.004, Mino: F_(1, 14)_ = 1.098, p = 0.312, Reminder x Mino: F_(1, 14)_ = 1.737, p = 0.209). Results are the mean ± SEM. n = 8 per group. * p < 0.05, ** p < 0.01, *** p < 0.001 compared between LTM tests; # < 0.05, ## p < 0.01 compared between groups.

In female rats conditioned using a 0.7mA that produced a robust long-lasting fear memory, minocycline administration had no effect on the retrieval, extinction or reinstatement by a reminder US after extinction of remote CFM as evidenced by no differences in freezing behaviour between the PBS or minocycline groups at any point during the experiment (Fig. 4C).

These data show that a 10 min exposure to the conditioning context can extinguish remote memory established for 25 days in both male and female rats. There was however a differential effect of minocycline treatment in males and females exemplified by higher conditioned freezing responses compared to controls in males notwithstanding evidence of extinction, suggesting minocycline enhances the reconsolidation of remote memory in males only.

## Discussion

We show that the acute administration of minocycline has profound effects on the expression of hippocampal-dependent contextual fear memory (CFM) in rats. The effect of minocycline was influenced by the sex of the rats, the age of the memory, and the conditions of memory recall associated with its administration. In males, minocycline given 48 hours after contextual fear conditioning (CFC), in the absence of a memory retrieval test, increased fear memory expression in subsequent tests for up to 23 days. However, when administered prior to retrieval, this potentiating effect was lost, and extinction could be enhanced, resulting in weaker expression of the fear memory. The enhancement of extinction by minocycline was dependent on the age of the fear memory and was limited to recently formed memories. Indeed, administering minocycline before retrieval of a well-established, 28-day-old memory actually increased the fear response to conditioned contexts by enhancing reconsolidation of the CFM in males. In contrast, in females trained under conditions that produced similar levels of fear memory expression to males, minocycline increased the behavioural expression of recent fear memory, regardless of whether the memory was retrieved following administration. It had no effect on extinction. These data suggest that using minocycline to ameliorate fear memories associated with psychiatric disorders may be limited by sex, the presence of extinction training, and the age of the fear memory. Specifically, moderation of fear memory was restricted to males undergoing extinction of recently acquired fear memories. Furthermore, minocycline treatment outside of these specific conditions may risk exacerbating fear memory expression.

In males, two days after fear conditioning, a single administration of minocycline before retrieval enhanced the behavioural extinction of contextual fear responses. This potentiating effect of minocycline on extinction was revealed under stronger contextual fear conditioning (CFC) conditions, which produced higher levels of conditioned freezing behaviour, and when repeated, prolonged (10-minute) re-exposures to the training context were provided. Under these conditions, minocycline resulted in lower freezing responses compared to matched controls at later time points. The non-reinforced retrieval of fully established memory can initiate the loss of memory-associated behaviour either through extinction[33], a new learning process in which conditioned environmental stimuli acquire new predictive properties (a CS–no US association; [33, 34] ) which competes with the original memory (CS– US association) for control over the behavioural response, or alternatively through the destabilization and restabilization of the original memory (CS–US) via reconsolidation [51, 52]. The recovery of freezing responses after a reminder US in our experiments, observed in both minocycline-treated and control groups, suggests that minocycline primarily enhanced extinction processes (context–no US learning), rather than blocking reconsolidation or causing a loss of the original contextual fear memory (context–US association), which would have been evident as a lack of recovery in freezing behaviour. This observation is consistent with extinction being favoured by the retrieval of stronger memories as well when there is a larger mismatch between the conditions of training and retrieval, including features such as the stimuli present and length and number of trials [53]. Furthermore, a direct effect of minocycline on reconsolidation processes on recently acquired CFM was not apparent; when minocycline was administered prior to a brief (2-minute) recall session, which typically favours engagement of reconsolidation, there was no observed difference in long-term memory expression compared to controls, measured up to 23 days later. Therefore, in males, minocycline appears to preferentially modulate extinction rather than reconsolidation of recently acquired fear memory.

The effect of minocycline in enhancing the extinction of recent CFM in males was broadly consistent with previous reports [32, 54]. Unlike these earlier studies, in which minocycline was administered chronically, potentially within the window for consolidation of the original fear memory, this study used a single dose administered outside that window and prior to extinction training. This provides more direct evidence for minocycline’s effect on extinction processes specifically engaged by CFM recall. There appeared to be no effect of minocycline on within-session extinction (i.e., during extinction training itself); rather, the drug influences fear memory expression during later recall. This suggests that, in males, minocycline most likely affects the consolidation of long-term extinction memory rather than its initial acquisition.

Minocycline in males increases fear memory expression when given without memory recall, contrasting its extinction-enhancing effects during retrieval. This suggests minocycline impairs natural (passive) forgetting of established fear memories [55, 56]. A single dose strengthened both strong (CFC using a single 2s 0.5mA US) and weak (trained with a 0.35 mA US) contextual fear memories lasting over 23 days, aligning with findings that minocycline prevents forgetting in male mice [31]. Thus, minocycline affects two memory processes: enhancing extinction (the formation of a context–no event association), when given before retrieval and reducing forgetting (preserving the context–US association) when administered without recall, resulting in nominally opposing effects on fear expression.

In contrast to the differential effects of acute minocycline on the expression of CFM observed in males, female rats showed a consistent enhancement of conditioned fear behaviour following minocycline administration two days after CFC, regardless of whether the drug was given in the absence or presence of memory retrieval (see Supplementary Fig. 1 for additional analysis of the enhancing of minocycline on retrieved CFM in females). Although higher US intensities were used during CFC in females to produce a robust and enduring CFM comparable to those of males, females nevertheless exhibited extinction and forgetting similar to males. However, unlike males, there was no evidence that minocycline modulated extinction or forgetting processes in females. The increased conditioned responses after minocycline without retrieval may reflect a nonspecific rise in fear and anxiety-like behaviours, rather than effects on memory processes. A recent meta-analysis showed that minocycline reduces anxiety-like behaviour in rodents, but only in those previously exposed to chronic stress or immune challenges, mainly studied in males [57]. There is a link between stress- induced anxiety response and neuroinflammation[58–60]. Together these data suggests that neuroinflammatory status of the animal may be a factor in determining the expression of fear and anxiety like behaviours in response to minocycline (see below for further discussion).

The effects of acute minocycline administration were critically dependent on the temporal age of the fear memory. In males, administration within two days of learning facilitated extinction training, leading to a reduction in fear-related behaviour. In contrast, delayed administration resulted in enhanced fear expression, even following extinction training. The inability of a reminder to reinstate fear behaviour after extinction suggests that minocycline may strengthen the original conditioned stimulus–unconditioned stimulus (CS– US) association via reconsolidation. The capacity of minocycline to differentially modulate extinction or reconsolidation likely relates to the changes in the conditions of retrieval required to engage either of the two processes (‘’boundary” conditions) and which are sensitive to memory age [53]. In females, the facilitatory effect of minocycline on fear expression was observed for recently acquired, but not remote, memories. Collectively, these findings indicate that the effects of minocycline on fear memory are contingent on both memory age and sex: in males, shifting from enhancing extinction and attenuating cued fear expression to promoting reconsolidation and amplifying fear responses, whereas in females, the capacity of minocycline to enhance fear expression was restricted to recent memories.

This study did not investigate the specific targets or mechanisms by which minocycline affects fear memory processing. However, minocycline is known to have anti-inflammatory and neuroprotective effects by regulating microglial activation [24, 61]. Microglia modulate neuronal activity and remodel neural circuits via cytokines, neurotrophic factors, and synaptic elimination—processes essential for learning and memory [31, 62, 63]. These disparate functions reflect the dynamic, plastic and possibly transitory morphological, transcriptomic and proteomic states of microglia [64–66]. Minocycline reduces complement-dependent synaptic elimination needed for forgetting contextual fear [31] and inhibits microglial TNF-α, promoting extinction [32]. Thus, minocycline’s effects in male rats, limiting forgetting while enhancing extinction, may involve modulation of microglial functions. These opposing effects may stem from changes in microglial activation triggered by neuronal activity during retrieval. Evidence suggests distinct microglial states underlie fear conditioning and extinction [67, 68], supporting the idea that microglia contribute to forgetting and extinction via separate mechanisms, warranting further study.

There is increasing evidence of sex differences in fear learning, memory, and extinction, driven by overlapping yet distinct molecular and neural mechanisms [69–72]. Differences also extend to memory retention, including extinction memory [73, 74]. Microglial activity may underlie these variations, as microglia differ by sex in density, distribution, molecular profiles, and function [75–79]. Moreover, microglial responses to stress, which affect fear and anxiety-like behaviours, are also sex-specific [80–82]. For example, microglial activation impairs fear extinction after chronic stress in males but not females [83]. Minocycline has been shown to reduce anxiety-like behaviour in both sexes, possibly through sex-specific mechanisms, such as reduced microglial phagocytosis in females [84, 85]. These findings suggest that minocycline’s differing effects on fear and anxiety-related behaviours may stem from sex-specific microglial states and responses to stress or stress associated memory retrieval.

In summary, our findings show that the effects of acute systemic minocycline on fear memory and anxiety-like behaviours in rats depend on sex, memory activation (retrieved or not), and memory age. In males, minocycline enhanced extinction of recently acquired fear when given before retrieval but increased fear expression when administered without recall, likely by disrupting passive forgetting. This effect was absent for older (28-day-old) memories where retrieval instead appeared to allow minocycline to strengthen the memory via reconsolidation. In females, minocycline increased fear expression memory regardless of retrieval shortly after training without affecting extinction or forgetting, suggesting a link to heightened anxiety-like behaviour. These results suggest that minocycline’s impact may depend on sex- and context-specific microglial states. Therapeutically, this implies restricted and condition-specific usefulness—for example, as an adjunct to exposure therapy for recent fear memories in men, while raising caution about potential adverse effects in other contexts. Such variability may help explain inconsistent clinical trial outcomes where these factors were not taken into account.

## Acknowledgements

This research was funded by the Hodge Centre for Translational Neuroscience at Cardiff University with further support from legacy funds.

---

**Supplementary Fig 1.**
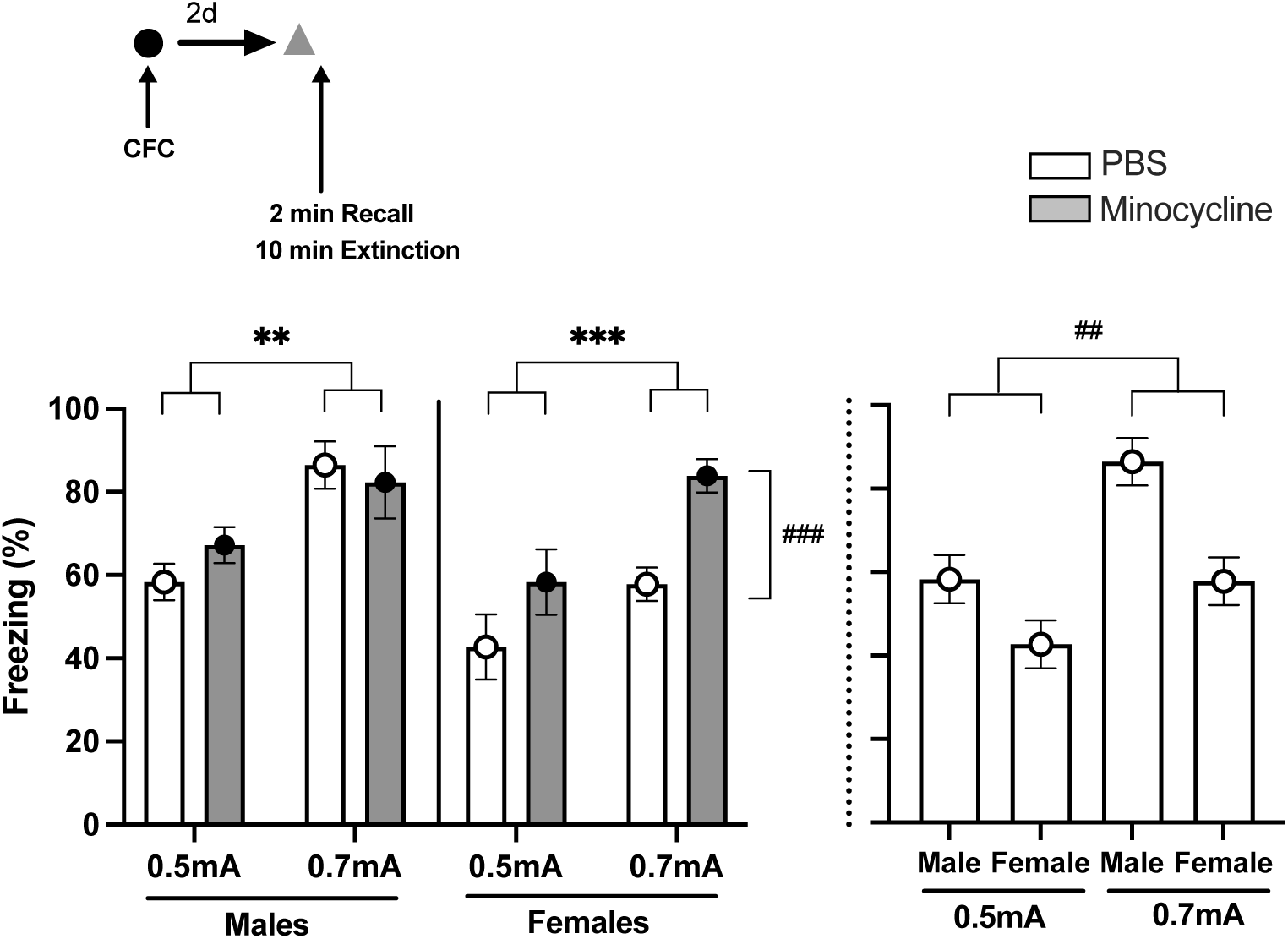
CFM is correlated to US intensity during CFC in males and females but minocycline enhances the expression of CFM only in females. Top: Schematic of the CFC and CFM behavioural protocol. CFC was established using a single 2s scrambled footshock US (2s, 0.5 mA or 2s, 0.7 mA) given 2min into a 3 min exposure to a novel context. Arrowhead shows the administration of minocycline (40mg/kg, i.p.) or PBS 30 min prior to 2min CFM recall trial. *Left*: Higher US’s evoked greater freezing behaviour with CFM retrieval 2 days after CFC in both sexes but minocycline increased freezing behaviour in females but not males (Males: US: F_(1, 76)_ = 7.879, p = 0.006, Mino: F_(1, 76)_ = 0.093, p = 0.762, US x Mino: F_(1, 76)_ = 0.715, p = 0.401. Females: US: F_(1, 52)_ = 11.853, p = 0.001, Mino: F_(1, 52)_ = 12.469, p < 0.001, US x Mino: F_(1, 52)_ = 0.779, p = 0.381)). *Right*: Noting the caveat that the males and females were conditioned and tested at different times, a planned comparison of data contained within lefthand panel suggested that in non-drug administered (control) rats, females showed lower levels of freezing behaviour than control males (F(_(1, 68)_ = 9.366, p = 0.003). The data represent the analysis of the combined freezing behaviour of all rats shown in Figs. 1C, 1D, 1F, 1G, 2A, 2B, 3C and 3D during the 1^st^ 2 min of the LTM test after minocycline or vehicle injection. Results are the mean ± SEM. Males (0.5mA: PBS: n = 32, minocycline: n = 32; 0.7mA: PBS: n = 8, minocycline: n = 8), Females (0.5mA: PBS: n = 16, minocycline: n = 8; 0.7mA: PBS: n = 16, minocycline: n = 16). **p < 0.01.

**Supplementary Table 1.**
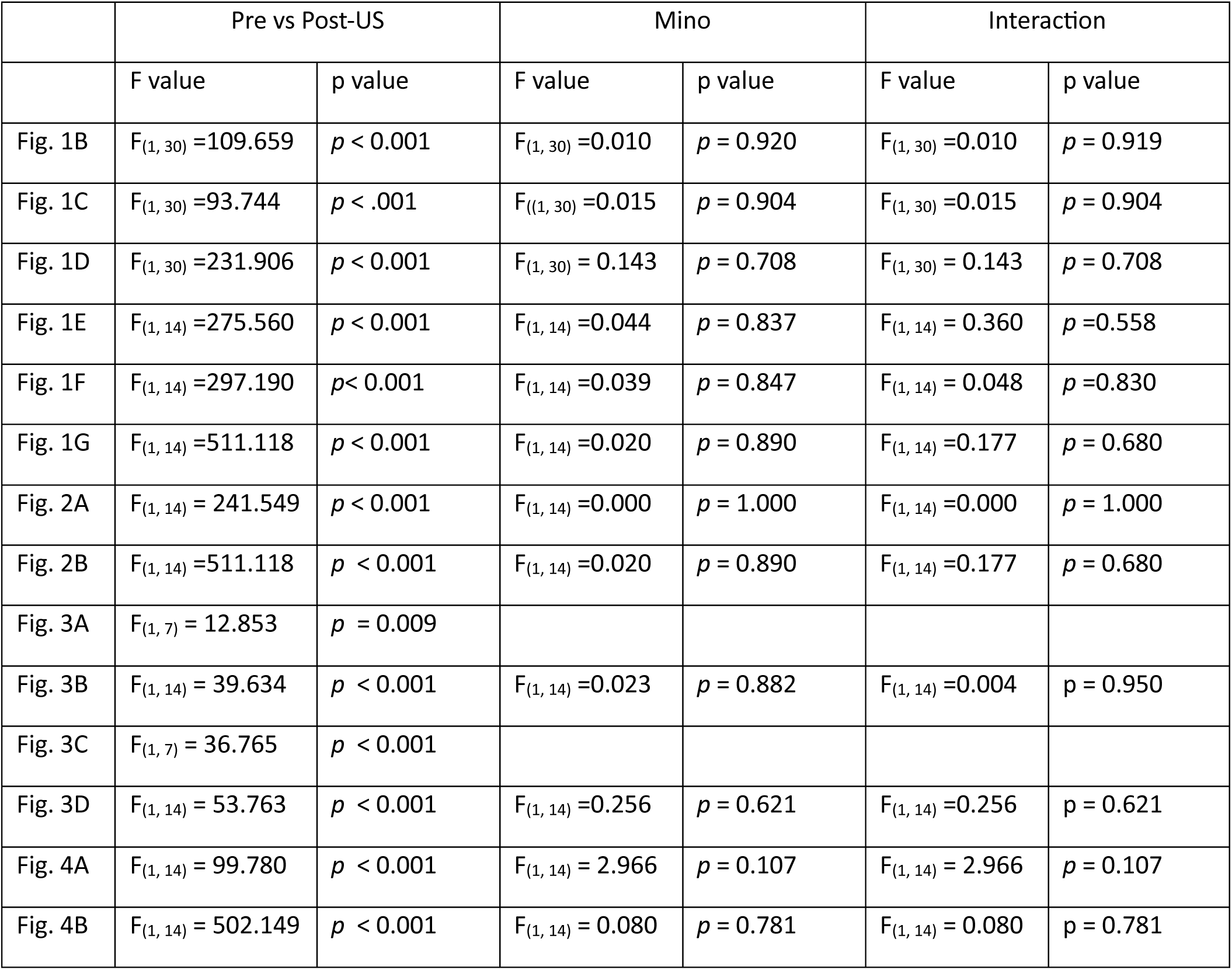
ANOVA reports of the freezing behaviour prior to and post US (footshock) during CFC for each experiment reported in Figs. 1-4. Footshock increased freezing responses in rats during CFC training and there were no group differences in freezing behaviour prior to subsequent systemic i.p. injections or behavioural manipulations.

