## Supplemental Table 1 for "Sex Dependent Effects of Minocycline on Contextual Fear Memory"

**Supplementary Table 1**

ANOVA reports of the freezing behaviour prior to and post US (footshock) during CFC for each experiment reported in Figs. 1-4. Footshock increased freezing responses in rats during CFC training and there were no group differences in freezing behaviour prior to subsequent systemic i.p. injections or behavioural manipulations.

| | Pre vs Post-US | | Mino | | Interaction | |
|---|---|---|---|---|---|---|
| | F value | p value | F value | p value | F value | p value |
| Fig. 1B | $F_{(1, 30)}$ =109.659 | $p < 0.001$ | $F_{(1, 30)}$ =0.010 | $p = 0.920$ | $F_{(1, 30)}$ =0.010 | $p = 0.919$ |
| Fig. 1C | $F_{(1, 30)}$ =93.744 | $p < .001$ | $F_{((1, 30)}$ =0.015 | $p = 0.904$ | $F_{(1, 30)}$ =0.015 | $p = 0.904$ |
| Fig. 1D | $F_{(1, 30)}$ =231.906 | $p < 0.001$ | $F_{(1, 30)}$ = 0.143 | $p = 0.708$ | $F_{(1, 30)}$ = 0.143 | $p = 0.708$ |
| Fig. 1E | $F_{(1, 14)}$ =275.560 | $p < 0.001$ | $F_{(1, 14)}$ =0.044 | $p = 0.837$ | $F_{(1, 14)}$ = 0.360 | $p =0.558$ |
| Fig. 1F | $F_{(1, 14)}$ =297.190 | $p< 0.001$ | $F_{(1, 14)}$ =0.039 | $p = 0.847$ | $F_{(1, 14)}$ = 0.048 | $p =0.830$ |
| Fig. 1G | $F_{(1, 14)}$ =511.118 | $p < 0.001$ | $F_{(1, 14)}$ =0.020 | $p = 0.890$ | $F_{(1, 14)}$ =0.177 | $p = 0.680$ |
| Fig. 2A | $F_{(1, 14)}$ = 241.549 | $p < 0.001$ | $F_{(1, 14)}$ =0.000 | $p = 1.000$ | $F_{(1, 14)}$ =0.000 | $p = 1.000$ |
| Fig. 2B | $F_{(1, 14)}$ =511.118 | $p < 0.001$ | $F_{(1, 14)}$ =0.020 | $p = 0.890$ | $F_{(1, 14)}$ =0.177 | $p = 0.680$ |
| Fig. 3A | $F_{(1, 7)}$ = 12.853 | $p = 0.009$ | | | | |
| Fig. 3B | $F_{(1, 14)}$ = 39.634 | $p < 0.001$ | $F_{(1, 14)}$ =0.023 | $p = 0.882$ | $F_{(1, 14)}$ =0.004 | p = 0.950 |
| Fig. 3C | $F_{(1, 7)}$ = 36.765 | $p < 0.001$ | | | | |
| Fig. 3D | $F_{(1, 14)}$ = 53.763 | $p < 0.001$ | $F_{(1, 14)}$ =0.256 | $p = 0.621$ | $F_{(1, 14)}$ =0.256 | p = 0.621 |
| Fig. 4A | $F_{(1, 14)}$ = 99.780 | $p < 0.001$ | $F_{(1, 14)}$ = 2.966 | $p = 0.107$ | $F_{(1, 14)}$ = 2.966 | $p = 0.107$ |
| Fig. 4B | $F_{(1, 14)}$ = 502.149 | $p < 0.001$ | $F_{(1, 14)}$ = 0.080 | $p = 0.781$ | $F_{(1, 14)}$ = 0.080 | p = 0.781 |
