## Supplementary figures and images for "Sex Dependent Effects of Minocycline on Contextual Fear Memory"

### Supplemental Figure 1

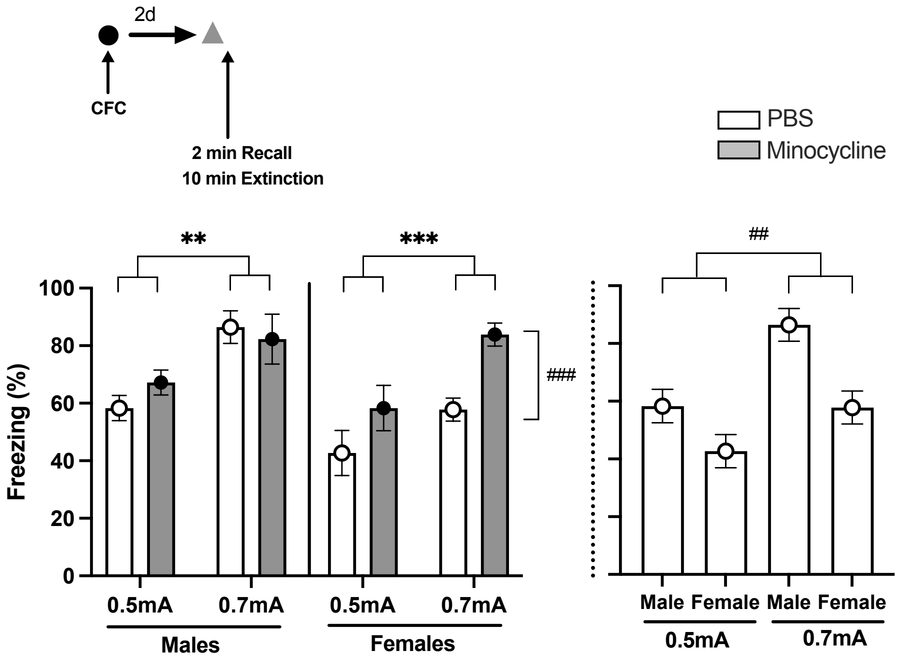
